# Barking up the wrong tree: the importance of morphology in plant molecular phylogenetic studies

**DOI:** 10.1101/2023.01.30.526223

**Authors:** Rafael Felipe de Almeida, Martin Cheek, Marco O.O. Pellegrini, Isa L. de Morais, Rosangela Simão-Bianchini, Pantamith Rattanakrajang, Ana Rita G. Simões

**Author notes:** Corresponding author: Rafael Felipe de Almeida –.

## Abstract

**Background and aims:** *Keraunea* is a genus recently described in Convolvulaceae, though it has sat uncomfortably in this family. A recent molecular phylogenetic study suggests that its two morphologically almost identical species actually belong to different families, Malpighiaceae (Superrosids) and Ehretiaceae (Superasterids), although with little-to-no morphological evidence to support it.

**Material and methods:** Sequences of *matK, rbcL*, and ITS for all the 77 currently accepted genera of Malpighiaceae, *K. brasiliensis* and Elatinaceae (outgroup) were compiled from Genbank and analysed with Maximum Likelihood and Bayesian Inference criteria for nuclear, plastid and combined datasets. Additional database and herbarium studies were performed to locate and analyse all duplicates of the holotype of *K. brasiliensis* to check for misidentified or contaminated materials.

**Key results:** Our examination of expanded DNA datasets and herbarium sheets of all *K. brasiliensis* isotypes revealed that an error in tissue sampling was, in fact, what led to this species being placed in Malpighiaceae. Kew’s isotype had a leaf of Malpighiaceae (likely from *Mascagnia cordifolia*) stored in the fragment capsule, which was unfortunately sampled and sequenced instead of the actual leaves of *K. brasiliensis*.

**Conclusions:** DNA sequences can be helpful in classifying taxa when morphology is conflicting or of a doubtful interpretation, with molecular phylogenetic placement becoming a popular tool that potentially accelerates the discovery of systematic relationships. However, good knowledge of plant morphology is essential for formulating the phylogenetic hypotheses to be tested and for a critical re-interpretation of the results in the context of biological information of the species or families. Thus, these techniques are, much like any others, prone to methodological errors. We highlight the crucial need to observe plant morphology alongside molecular phylogenetic results, particularly when the new hypotheses are in disagreement with the existing classification and at risk of incurring gross taxonomic mistakes.

## Introduction

Plant phylogenetic studies have relied on herbarium collections for DNA extraction for the past four decades due to the easy access to a wealth of specimens that could, otherwise, only be obtained via costly or difficult fieldwork, such as: 1. specimens from a wide range of geographical locations (e.g., different continents); 2. scarce specimens (e.g., type specimens), 3. threatened or extinct species; and 4. geographically restricted populations (Shepherd and Perrie 2014; Bieker and Martin 2018). Since the DNA in mounted herbarium specimens’ decays six times faster than in bone, it is usually available in small amounts and highly degraded into short DNA fragments (Weiß et al. 2016; Bieker and Martin 2018). Although some herbarium specimens are simply too degraded to be used in traditional sequencing methods (i.e., Sanger sequencing), Next Generation Sequencing methods, which enrich DNA extracts with extremely short (40–00 bp) DNA fragments, pushed the boundaries of what could be sequenced from very low-quality genetic material (Bieker and Martin 2018). These new advances in DNA sequencing from herbarium specimens have been reflected in the last decade of published phylogenetic studies, with an increase of 50% in the number of studies mainly or solely relying on herbarium specimens for DNA extraction (Bieker and Martin 2018).

A common issue with using herbarium samples for DNA extraction is contamination and misidentification, usually occurring during laboratory procedures for DNA obtention (Wang 2018) or originating from the biological collections themselves when specimens are not identified by taxonomic experts. Several new phylogenomic studies have been proposing new statistical methods to detect and remove contaminants from animal (Weissensteiner et al. 2021; Owen et al. 2022), bacterial (Pightling et al. 2019), or environmental (Sepulveda et al. 2021) phylogenomic datasets. However, before these recent efforts to identify contaminants in phylogenomic datasets were developed, previous phylogenetic studies that included contaminated or misidentified sequences generated erroneous phylogenetic trees in different groups of the plant tree of life (e.g., Apiaceae – Downie et al. 2010; Betulaceae – Wang et al. 2016; Juncaceae – Elliot et al. 2023; Laurales – Smith and Brown 2018; Menispermaceae – Ortiz et al. 2007; Rubiaceae – McCartha et al. 2019; and Tofieldiaceae – Chen et al. 2013). The impact on subsequent secondary analyses of these genetic data is massive, such as molecular dating estimates, biogeographic inferences, or systematic studies re-classifying organisms solely based on the contaminant sequences (Wang et al. 2014). Nonetheless, to the best of our knowledge, no study to date has addressed the impact of unskilful tissue sampling (i.e., the lack of taxonomic expertise or morphological knowledge of the plant groups) on herbarium samples used for DNA extraction in plant molecular systematics.

The genus *Keraunea* was first published by Cheek and Simão-Bianchini (2013) based on a single species endemic to Brazil, *K. brasiliensis* Cheek & Sim.-Bianch. At the time, it was proposed to belong to the family Convolvulaceae, based on the presence of a superior ovary with two carpels, bifid stigma, gamopetalous corolla, epipetalous stamens, climbing habit, and alternate, exstipulate, pinnately-nerved, simple, and entire leaves. It also presented an unusual fruit, with much-enlarged bracts, adnate to the pedicel (Figure 1), which is characteristic of *Neuropeltis* Wall., in tribe Poraneae (*sensu* Staples and Brummitt 2007). Another three genera of Convolvulaceae in this tribe, *Calycobolus* Willd. ex Schult., *Dipteropeltis* Hallier f. and *Rapona* Baill., present superficially similar wind-dispersed analogous structures, although, in these genera, they represent enlarged sepals instead of bracts embracing the fruit. The presence of a single style, rather than a bifid style, as is characteristic of *Neuropeltis* and allied genera, led Cheek and Simão-Bianchini (2013) to describe it as a separate genus, given the taxonomic significance of style characters at the generic level in Convolvulaceae. However, the conflict between the putative morphological similarities between *Keraunea* and *Neuropeltis*, and the genus’ clear distinctiveness from the other members of Poraneae, especially regarding style shape, resulted in it uncomfortably sitting within Convolvulaceae since its description.

**Figure 1.**
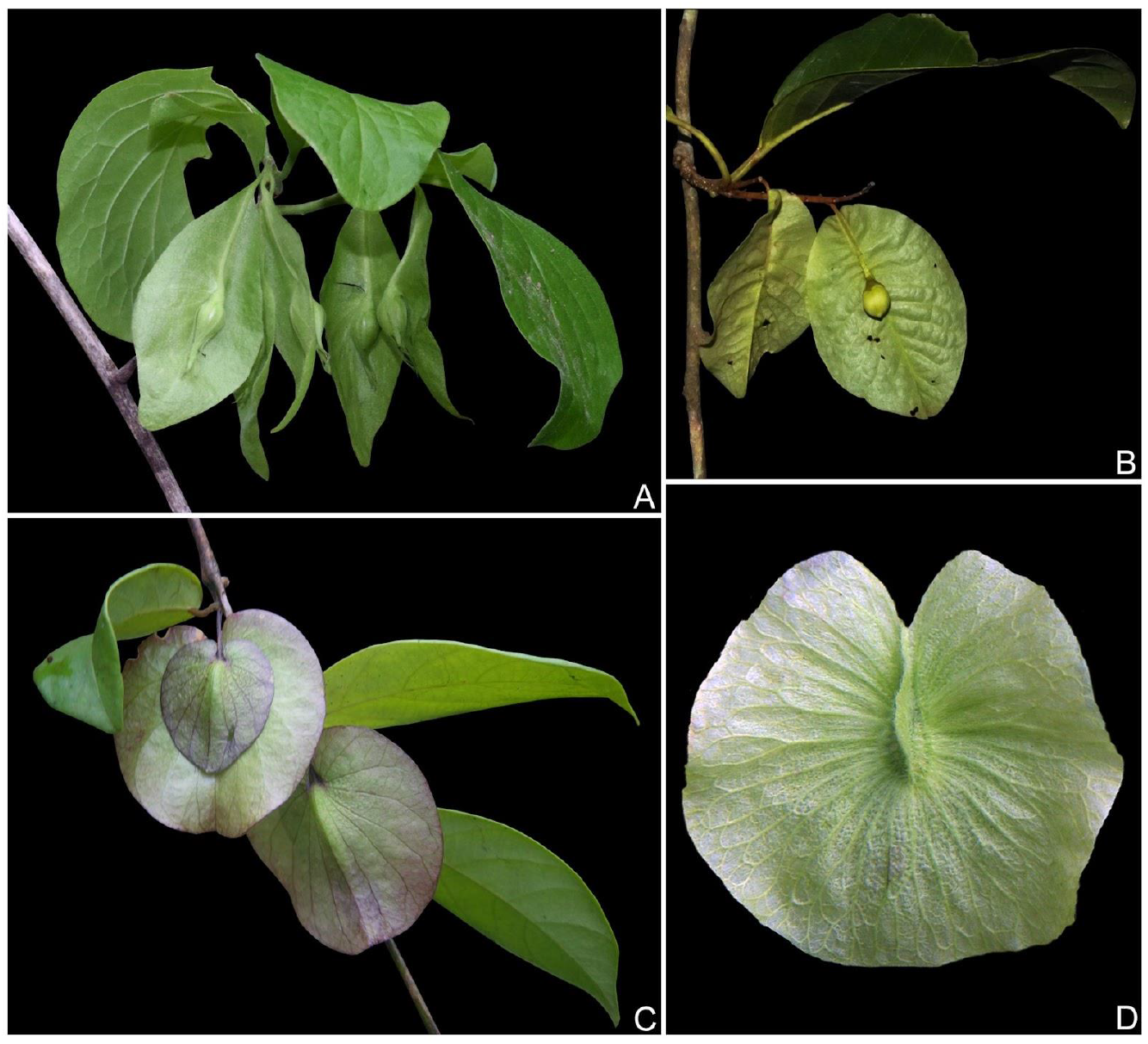
Field photographs of the fruits of (A) *Keraunea brasiliensis* Cheek & Simão-Bianchini (*incertae sedis*), photo by Domingos Cardoso; (B) *Neuropeltis racemosa* Wall. (Convolvulaceae), photo by P. Rattanakrajang; (C) *Calycobolus campanulatus* (K.Schum. ex Hallier f.) Heine (Convolvulaceae), photo by O. Lachenaud, and (D) *Mascagnia cordifolia* (A.Juss.) Griseb. (Malpighiaceae), photo by M.O.O. Pellegrini.

Only a few months after the genus’ description, a second species with comparatively minor morphological differences – such as indumentum density, inflorescence structure, the length of the calyx and corolla lobes, and corolla length – was added to *Keraunea, K. capixaba* Lombardi (Lombardi 2014). The description of the genus and both recognised species was solely based on macromorphological characters, with the molecular phylogenetic relationships of the species and the genus to the rest of Convolvulaceae remaining unconfirmed. A recent large-scale phylogenetic study of the family, using nuclear genomic data with target capture techniques (Simões et al. 2022), proposed that, alas, the genus did not belong to Convolvulaceae. However, no discussion on morphological characters supporting this placement was discussed, nor was an alternative family classification proposed, leaving *Keraunea* as “*incertae sedis*”.

Another recently published molecular phylogenetic study (Muñoz-Rodríguez et al. 2022), based on Sanger sequencing data, proposed to address the family placement of *Keraunea* by phylogenetic analysis of plastid (*matK, rbcL*) and nuclear (ITS) regions. Both species of the genus (i.e., *K. brasiliensis* and *K. capixaba*) were sampled, and sequences of a wide range of taxa across the family Convolvulaceae were added to the analysis. The phylogenetic analyses reinstated that the genus was, indeed, placed outside of Convolvulaceae, supporting Simões et al. (2022). Thus, Muñoz-Rodríguez et al. (2022) tested the monophyly of *Keraunea* for the first time, with the genus being deemed polyphyletic based only on molecular data and without providing morphological support. Furthermore, the authors state that the type species (i.e., *K. brasiliensis*) actually belongs to Malpighiaceae (Malpighiales, Superrosids) “despite several morphological anomalies”, while the second species (i.e., *K. capixaba*) belongs to Ehretiaceae (Boraginales, Superasterids). The authors also state that the isotype of *K. brasiliensis* (Passos 5263) was placed in Malpighiaceae, while one of the paratypes of the same species (Lombardi 1819) was placed in Ehretiaceae alongside *K. capixaba*. Consequently, these results raised serious suspicions of methodological issues when sampling Passos 5263. However, Muñoz-Rodríguez et al. (2022) conclude that “[their] molecular results strongly suggest Passos 5263 belongs to Malpighiaceae, most likely within *Mascagnia”* but refrain from proposing any taxonomic changes at that time.

We have found these results difficult to reconcile with the existing taxonomic knowledge of these plant groups, especially considering how evolutionarily distant Malpighiaceae and Ehretiaceae are and how both species of *Keraunea* are remarkably morphologically similar to each other. As a result, we have set ourselves to provide a more satisfactory explanation for these curious results with meticulous analyses of all available evidence. In this study, we test the placement of Passos 5263 (“*Keraunea brasiliensis*”) in Malpighiaceae through a set of rigorous and comprehensive phylogenetic and herbarium analyses of the same specimens and expanded DNA datasets as Muñoz-Rodríguez et al. (2022), shedding new light on this taxonomic conundrum. In future studies, we will discuss in depth the issues of the proposed placement of *K. capixaba* in Ehretiaceae, using expanded sampling and more advanced molecular techniques, not addressed here due to this work being ongoing and of greater underlying complexity.

## Material and Methods

### Phylogenetics

Muñoz-Rodríguez et al. (2022) used an outdated generic sampling for Malpighiaceae based on Davis and Anderson (2010). Since then, several phylogenetic and taxonomic studies have been published, including the synonymy of several genera and the publication of two new genera for Malpighiaceae (Davis et al. 2020, Almeida and van den Berg 2021). In order to accurately test the phylogenetic placement of *Keraunea brasiliensis* within Malpighiaceae, we downloaded sequences of the markers *matK*, *rbcL*, and ITS stored on Genbank for all 77 currently accepted genera of Malpighiaceae according to POWO (2023) and the sequences of *K. brasiliensis* (Passos 5263) presented as supplementary material by Muñoz-Rodríguez et al. (2022). Sequences of *Elatine* L. and *Bergia* L. (Elatinaceae), the sister group of Malpighiaceae (Cai et al. 2016; Davis and Chase 2004), were also used to root our analysis. Additionally, we performed a secondary analysis to test the placement of *K. brasiliensis* within the genus *Mascagnia* (Bertero ex DC.) Bertero, including 18 accepted species (out of 58) with available sequences on Genbank, four species of *Amorimia* W.R.Anderson as the outgroup, and *Ectopopterys* W.R.Anderson to root the analysis.

Datasets were compiled, for each marker, using the program Geneious (Kearse et al. 2012) and aligned using Muscle (Edgar 2004), with subsequent adjustments in the preliminary matrices by visual inspection. Separate and combined analyses of plastid, nuclear, and plastid + nuclear regions were performed using Bayesian Inference (BI) and Maximum Likelihood (ML) criteria for phylogenetic reconstruction. Both model-based methods were conducted with a mixed model (GTR+G+I) and unlinked parameters selected using J Modeltest 2 (Darriba et al. 2012), MrBayes 3.1.2 (Ronquist and Huelsenbeck 2003) and raxmlGUI2 (Edler et al. 2021). For the BI, the Markov Chain Monte Carlo (MCMC) was run using two simultaneous independent runs with four chains each (one cold and three heated), saving one tree every 1,000 generations for a total of ten million generations. We excluded 20% of retained trees as “burn in” and checked for a stationary phase of likelihood, checking for ESS values higher than 200 for all parameters on Tracer 1.6 (Rambaut et al. 2014). The posterior probabilities (PP) of clades were based on the majority rule consensus, using the stored trees, and calculated with MrBayes 3.1.2 (Ronquist and Huelsenbeck 2003). Support values are presented on branches, with bootstrap shown before posterior probabilities values.

### Herbarium studies

Images of the holotype and all isotypes of *Keraunea brasiliensis* (Passos 5263) were searched in online specimen databases, such as GBIF (https://www.gbif.org), JSTOR (https://plants.jstor.org), Jabot (http://jabot.jbrj.gov.br), Reflora (https://reflora.jbrj.gov.br), and speciesLink (https://specieslink.net). The Kew isotype (K000979156) was consulted in person at the Kew herbarium (Almeida, Simões and Cheek), and the Brazilian duplicates (ALCB, CEPEC, HRCB, HUEFS and SPF; acronyms according to Thiers, continuously updated) were accessed by Simão-Bianchini and local collaborators among the staff of the above-cited herbaria. At Kew herbarium, morphological details were photographed using a Leica S9i stereomicroscope with a coupled digital camera.

## Results

### Phylogenetics

The topologies of the nuclear, plastid and combined phylogenetic trees were found to be highly congruent. Hence, we have chosen to discuss the results in light of the combined analysis instead of the individual datasets (for additional information, see supplementary files). *Keraunea brasiliensis* (Passos 5263) was recovered with high support as nested within the genus *Mascagnia* in the Malpighioid clade by both the individual and combined analyses, using BI and ML inference criteria (Figure 2). The low support for most relationships within the Tetrapteroid clade is interpreted as the result of missing data between all three molecular datasets. Based on our secondary analysis, *K. brasiliensis* was recovered with high support as sister to *Mascagnia cordifolia* (A.Juss.) Griseb (Figure 3). Due to these curious results, we compared the DNA sequences of *K. brasiliensis* to *M. cordifolia* in the individual *matK* and *rbcL* alignments. The ITS dataset did not include a sequence of *M. cordifolia*. This careful analysis evidenced that these sequences had the exact same nucleotide composition (see supplementary files). Only the *rbcL* sequence of *K. brasiliensis* was shorter, with 600 bp long, instead of the ca. 1,400 bp long sequence for *M. cordifolia* and the remaining Malpighiaceae (see supplementary files).

**Figure 2.**
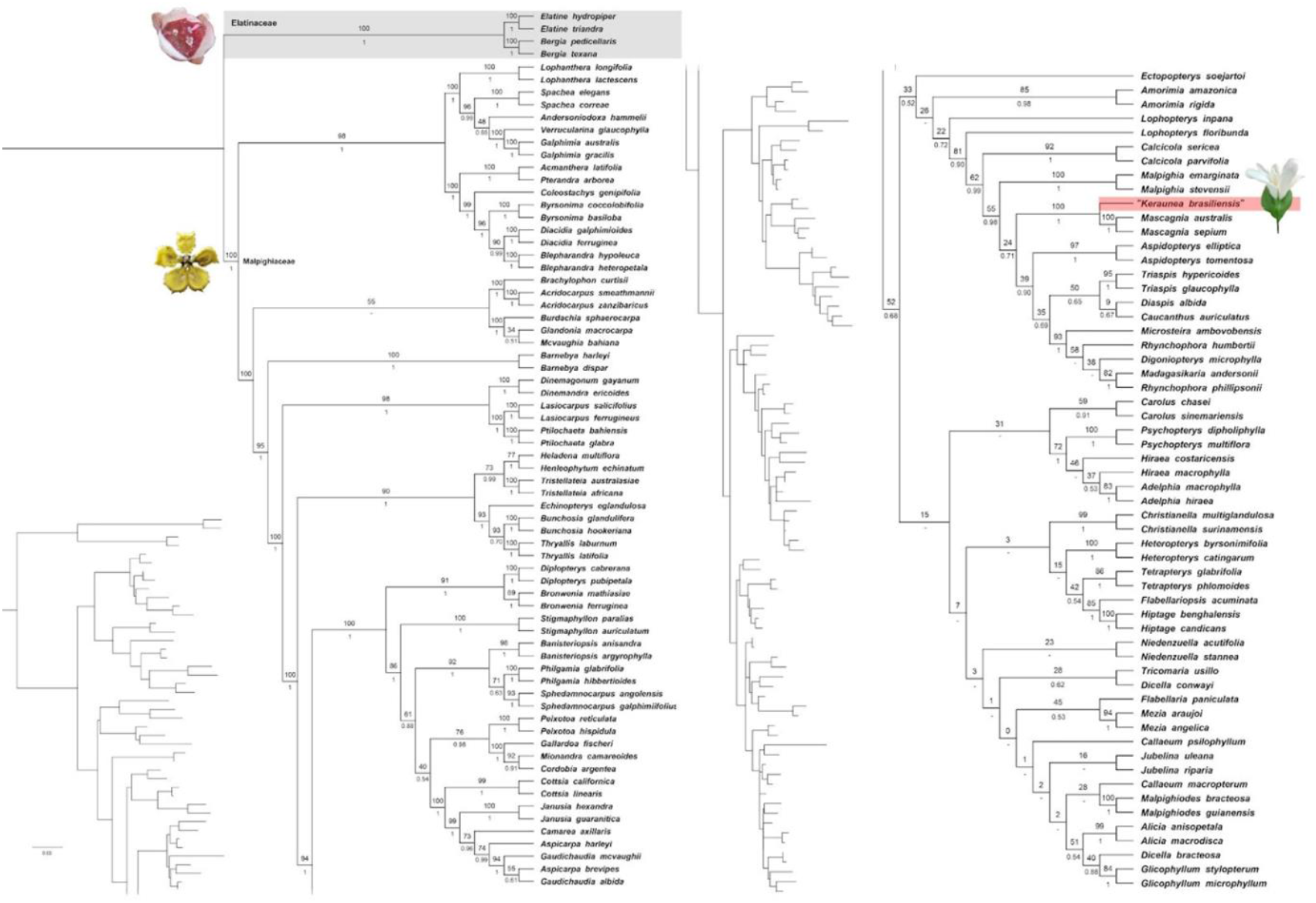
Consensus tree of the combined analysis based on the markers *matK, rbcL*, and ITS showing the phylogenetic placement of “*Keraunea brasiliensis* – Passos 5263” (highlighted in red) within Malpighiaceae, making both the family and the genus *Mascagnia* non-monophyletic. Elatinaceae (highlighted in light grey) represents the outgroup and the root of this analysis. Bootstrap values from the ML are shown above branches, and posterior probabilities from the BI are shown below branches. Trees on the left are presented for branch length visualisation. Photographs of *Elatine gratioloides* A.Cunn. by M. Hutchison, *Stigmaphyllon angustilobum* A.Juss. by R.F. de Almeida, and *Keraunea spp*. by G.S. Siqueira.

**Figure 3.**
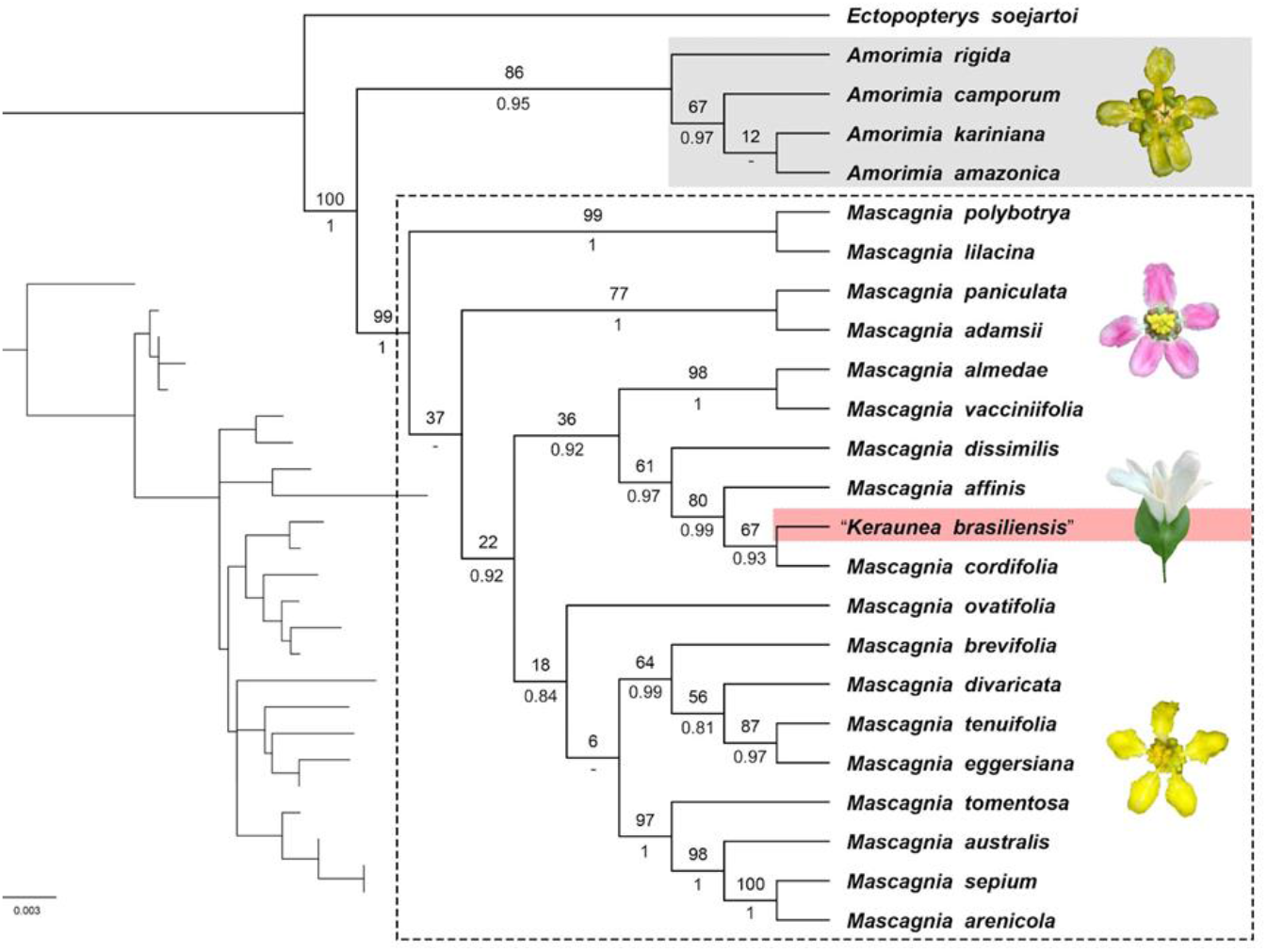
Consensus tree of the combined analysis based on the markers *matK* and *rbcL* showing the phylogenetic placement of *“Keraunea brasiliensis* - Passos 5263” (highlighted in red) within *Mascagnia. Amorimia* W.R.Anderson (highlighted in light grey) represents one of the outgroups, and *Ectopopterys* W.R.Anderson is the root of this analysis. Bootstrap values from the ML are shown above branches, and posterior probabilities from the BI are shown below branches. The tree on the left is shown for branch length visualisation. Photographs of *Amorimia ssp*. by Fabián Michelangeli, *Mascagnia cordifolia* by M.O.O. Pellegrini, and *Mascagnia australis* C.E.Anderson by C.F. Hall.

### Herbarium studies

Our molecular phylogenetic results agree with those of Muñoz-Rodríguez et al. (2022), placing their sequence of Passos 5263 in *Mascagnia*, very probably representing the species *Mascagnia cordifolia*, given the identical genetic sequences for the barcoding markers *matK* and *rbcL*. Hence, we moved on to questioning if there could have been, then, an issue with the source of the sequence, either by laboratory contamination or problems with the source of the samples. The first hypothesis of laboratorial contamination could be immediately discarded since Muñoz-Rodríguez et al. (2022) themselves stated that their sample of *K. brasiliensis* was re-sequenced by different laboratories, converging to identical sequences. This led us to investigate next if there could be an issue with the source of the sample. Hence, the solution to this taxonomic conundrum would rely on the analysis of the herbarium sheets sampled by Muñoz-Rodríguez et al. (2022).

Nonetheless, these authors do not explicitly state which duplicate of Passos 5263 was sequenced (from the six available types: ALCB, CEPEC, HRCB, HUEFS, K or SPF), although it is mentioned by the authors that only the K herbarium was visited in person. However, the K isotype (K000979156) does not present any annotation of having been sampled for DNA studies by the authors. In fact, this isotype only had a DNA sample slip dated from 2019 for unpublished molecular phylogenetic studies led by Kew’s in-house researcher Dr Tim Utteridge. Thus, we also looked at the possibility of the sequences of *K. brasiliensis* generated by Muñoz-Rodríguez et al. (2022) being from any of the Brazilian isotypes (i.e., ALCB037775, CEPEC00077827, HRCB38156, and HUEFS0028681) or the holotype (i.e., Passos 5263, SPF). The curators of the abovementioned herbaria were contacted, and it was confirmed that lead material for DNA sequencing had not been sent from these herbaria to this team of authors. Additionally, the database of the Brazilian federal government authority which regulates and authorises the use of any genetic material from Brazilian biological diversity in scientific studies (SisGen - Sistema Nacional de Gestão do Patrimônio Genético e do Conhecimento Tradicional Associado, https://sisgen.gov.br), was checked for records of authorisation having been granted to sequence the DNA of *K. brasiliensis* from any of the Brazilian herbaria in which all the type materials of *K. brasiliensis* were deposited. This search retrieved no results, further supporting that none of these materials had been sampled by foreign researchers or sequenced outside Brazil. Hence, we conclude that the sampled specimen would have, indeed, been the K sheet.

During the observation of the K isotype specimen, it was found that the plant was entirely glued to the sheet, and some detached leaves and fruits were stored in a paper capsule on the left lower side of the sheet (Figure 4). The fragments were likely the source of the sequenced sample, considering the difficulties in collecting leaf material from the glued specimen. Inside the paper capsule, we recognised that one of the leaves did not match the general morphology of the remaining *K. brasiliensis* leaves (Figure 4). Inspection under a hand lens and stereomicroscope revealed that this “distinct” leaf clearly presented V-shaped, 1-celled Malpighiaceous hairs instead of the unbranched, stout multi-celled hairs of the remaining leaves of *K. brasiliensis* (Figure 4). On this discovery, Simão-Bianchini and local collaborators also checked the herbarium sheets of the holotype and other Brazilian isotypes. Particular attention was given, if present, to the content of fragment capsules for the presence of possible foreign leaf material mixed up with true leaves of *Keraunea*. This investigation confirmed that all Brazilian specimens of Passos 5263, and their fragment capsules, contained only samples of *K. brasiliensis* and that none of them contained leaf fragments with V-shaped, 1-celled Malpighiaceous hairs. This led us to conclude that this sample mixture only pertained to the K isotype, further supporting our suspicions that Muñoz-Rodríguez et al. (2022) did indeed sample the K specimen.

**Figure 4.**
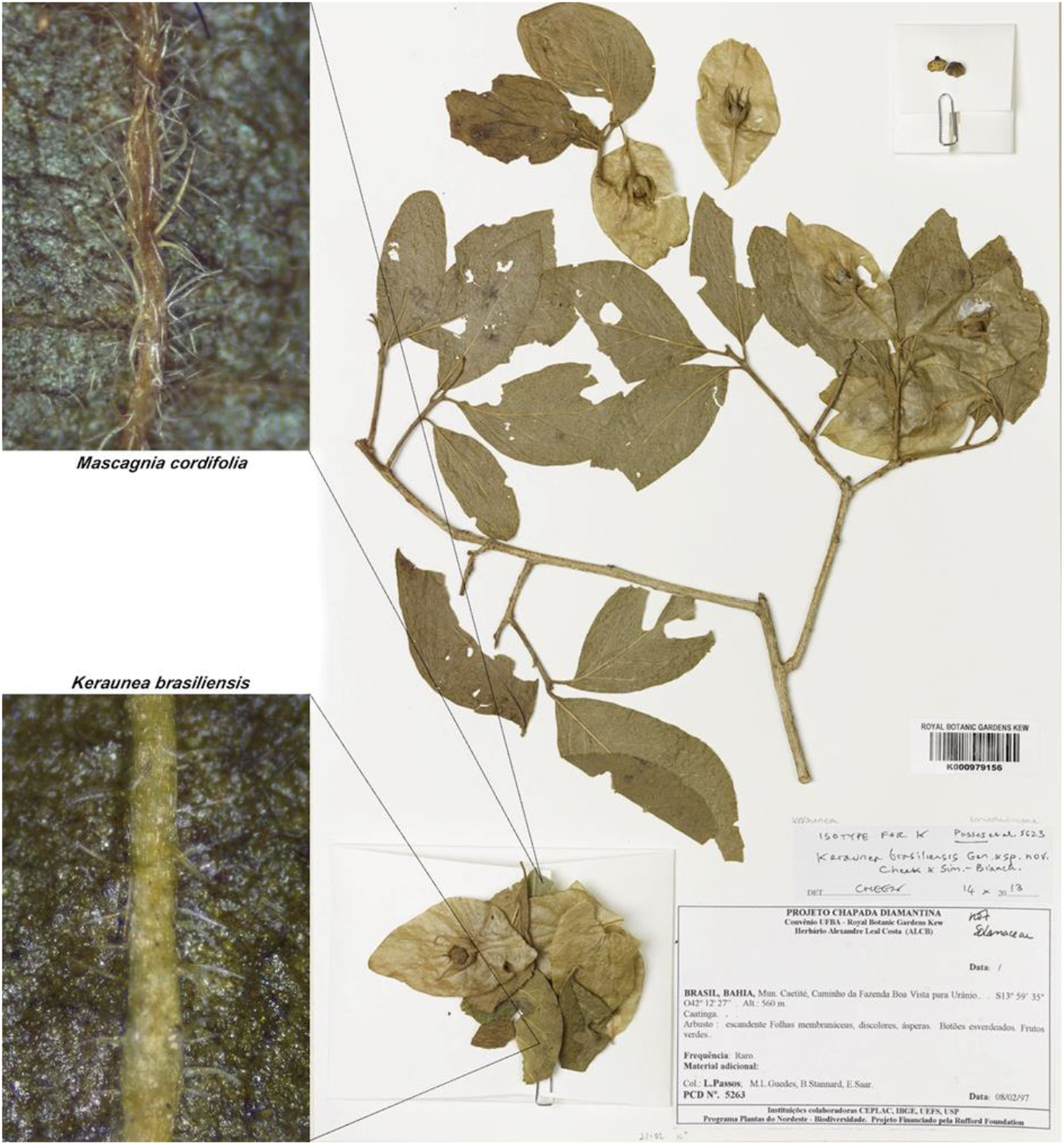
Photograph of the isotype of *Keraunea brasiliensis* (Passos 5263) deposited at RBG Kew’s herbarium showing the open fragment capsule storing leaves of *K. brasiliensis* (lower photographic detail showing unbranched and stout hairs) and a single leaf of *Mascagnia cordifolia* (upper photographic detail showing 2-branched hairs).

Thus, our critical re-analysis of the molecular and morphological evidence used by Muñoz-Rodríguez et al. (2022) allowed us to conclude that the phylogenetic placement of *K. brasiliensis* in Malpighiaceae can be explained by the contamination of the fragment capsule in the K isotype (Passos 5263, K000979156) by Malpighiaceae leaves and their fragments. It is possible that this contamination was accidentally introduced during fieldwork or during the processing of the specimens for herbarium incorporation when leaf material, most likely of *Mascagnia cordifolia*, was added to the capsule by mistake.

## Discussion

### Molecular phylogenetics of misidentifications and contaminants

It is not uncommon for molecular phylogenetic studies to evidence that genera, and sometimes species, are not monophyletic. In fact, the main contribution of molecular phylogenetic studies to plant systematics has been to challenge traditional classification systems based on morphological dogmas (Chase et al. 1993). The most famous case being the Dicotyledon/Monocotyledon traditional division of flowering plants, which led to an entirely different view of subdivisions of flowering plants after their first molecular phylogenetic studies demonstrated the non-monophyletic nature of the Dicotyledons (Chase et al. 1993). Nonetheless, to the best of our knowledge, *Keraunea* represents an unprecedented case in which its type species was proposed as non-monophyletic, with different samples of an isotype and a paratype being placed in distinct clades of Pentapetalae (i.e., Superasterids and Superrosids) without any morphological support.

When Muñoz-Rodríguez et al. (2022) proposed the non-monophyly of *K. brasiliensis*, these authors only re-sequenced their tissue samples in a different laboratory to corroborate their unusual results. In fact, this is only the first step towards troubleshooting phylogenetic incongruencies that are not corroborated by morphological evidence. The second and most important step is going back to the original sampled specimen to check for taxonomic misidentifications and/or contamination. Unfortunately, skipping this second step seems to be common practice in plant phylogenetic studies, with several Sanger DNA sequences published on Genbank for different groups of seed plants showing misidentification problems, as discussed by Smith and Brown (2018). These authors had to exclude several shorter gene regions (i.e., DNA barcode regions such as *matK* and *rbcL*) from their seed plant phylogenetic study due to misidentification or contamination that led to the incorrect placements of several of the analysed groups. Still, according to these authors, even if only a single sequence constitutes a misidentification or contamination, within a multiple loci analysis, it is enough to drive the incorrect placement of the taxon. Consequently, when conducting large-sized phylogenetic analyses, small percentages of bad data can dramatically inhibit accurate phylogenetic estimates (Smith and Brown 2018) and lead to much avoidable taxonomic upheaval, such as in the case of *Keraunea*.

### Systematics of Keraunea

The case of the *Keraunea brasiliensis* isotype being phylogenetically misplaced in Malpighiaceae would have been avoided if the sampler in the study of Muñoz-Rodríguez et al. (2022) had noticed the gross morphological differences in the leaves (i.e., different shape, texture, colouration, venation, base, margin and apex) or the presence of V-shaped, 1-celled Malpighiaceous hairs (Figure 4). Alternatively, in the face of the surprising molecular results, Muñoz-Rodríguez et al. (2022) should have critically re-analysed the sampled herbarium specimen while looking for the potential source of error. Even though most families of flowering plants are almost impossible to be reliably identified based only on vegetative characters, especially using leaf fragments, Malpighiaceae represents one of the few, and maybe one of the most well-known, exceptions. One of the most significant synapomorphies for this family, the indumentum of their vegetative structures, is always made of 1-celled hairs with two branches that can be T-, Y- or V-shaped, united by a well-developed or inconspicuous base (i.e., foot) (Almeida and Morais 2022).

This unique hair morphology was first described in Malpighiaceae and has since been referred to as Malpighiaceous hairs in taxonomic literature (Almeida and Morais 2022). However, Malpighiaceous hairs are not exclusive to Malpighiaceae, being also found in 23 unrelated families of flowering plants (i.e., Acanthaceae, Aizoaceae, Asteraceae, Boraginaceae, Brassicaceae, Burseraceae, Cannabaceae, Capparaceae, Combretaceae, Convolvulaceae, Cornaceae, Connaraceae, Ebenaceae, Escalloniaceae, Lythraceae, Myrtaceae, Sapindaceae, Sapotaceae, Thymelaeaceae, Verbenaceae, Vitaceae, Vochysiaceae, and Zygophyllaceae; Rao and Sarma 1992). Within Convolvulaceae, 2-branched hairs (i.e., Malpighiaceous hairs) are found in *Ipomoea* L. (Wood et al. 2016), *Evolvulus* L. (Silva and Simão-Bianchini 2014), *Cordisepalum* Verdc. (Staples 2006), *Dinetus* Buch.-Ham. ex D.Don (Staples 2006), *Duperreya* Gaudich. (Staples 2006), *Poranopsis* Roberty (Staples 2006), *Tridynamia* Gagnep. (Staples 2006), *Stylisma* Raf. (Myint 1966), *Jacquemontia* Choisy (Patel 2021), *Erycibe* Roxb. (Kochaiphat et al. 2021), and *Neuropeltis* Wall. (Breteler 2010).

*Keraunea* was proposed by Cheek and Simão-Bianchini (2013) to be close to *Neuropeltis*, a rather unusual Palaeotropical genus of Convolvulaceae with very enlarged bracts, partly fused to the pedicel, contrasting with another odd African genus, *Calycobolus*, in which the sepals are the organs which enlarge, instead of the bracts, forming a membranous structure, completely enclosing the fruit. The enlarged membranous bracts of *Keraunea* greatly resemble the membranous to subcoriaceous bracts present in *Neuropeltis* and *Neuropeltopsis* Ooststr., being one of the first morphological characters that suggested its original placement in Convolvulaceae. Nonetheless, molecular phylogenetic evidence did not corroborate the close relationship between *Keraunea* and *Neuropeltis* (Muñoz-Rodríguez et al. 2022; Simões et al. 2022), although sampling of *Neuropeltis* was limited to only one African species, *N. acuminata* (P.Beauv.) Benth.

Since *Neuropeltis* shows Malpighiaceous hairs in several species (Breteler 2010), one could argue that this type of hair is common in several genera of Convolvulaceae, being the main reason justifying the erroneous sampling by Muñoz-Rodríguez et al. (2022). Nonetheless, a minute but crucial detail distinguishes the hairs of Convolvulaceae from those of Malpighiaceae. In Convolvulaceae, the 2-branched hairs are always 2-celled and T-shaped, while in Malpighiaceae, the 2-branched hairs are always 1-celled and T-, Y- or V-shaped, such as in the mixed material from Passos 5263 (K000979156; Figure 4). Therefore, based solely on morphology, it can be confidently established that the mixed material from the isotype at Kew is of a Malpighiaceae species, most probably a specimen of *Mascagnia cordifolia*, which always shows V-shaped hairs such as those presented in Figure 4. Additionally, the hairs present in all leaves of the known specimens of *Keraunea brasiliensis* are always unbranched, multi-celled, and bulbous at the base, as described by Cheek and Simão-Bianchini (2013) and confirmed by us when re-examining the specimens (Figure 4). Until a broader molecular sampling of Convolvulaceae is possible, namely of *Keraunea*, *Neuropeltis* and even *Calycobolus*, any placement proposed based on molecular phylogenetics alone could be misleading and speculative.

As we here conclude, this was the case regarding the proposed inclusion of one of the isotypes of *K. brasiliensis* in Malpighiaceae, which resulted from a methodological blunder. For this reason, the family placement of *Keraunea* as a whole is currently being analysed with extreme caution, with expanded sampling that includes holotypes, isotypes, paratypes and more recent collections of both species, and in the context of multiple sources of evidence (i.e., morphological, ecological, palynological, anatomical and molecular). This is being done to ensure the most robust family placement hypotheses and avoid erroneous or speculative proposals that could unnecessarily disrupt the existing classifications of other plant families (e.g., Malpighiaceae and Ehretiaceae) and, as a result, cause the erroneous re-interpretation of character evolution or biogeographic processes in those plant groups. Thus, we here refute the hypothesis of *Keraunea brasiliensis* – or *Keraunea pro parte* – belonging to Malpighiaceae and expect to more rigorously address its correct family placement in future studies.

### Future directions

These problems highlight the need to adequately address how molecular phylogenetic studies should proceed with contaminated and/or misidentified sequences for large-sized phylogenetic analyses. Hence, taxonomic expertise is fundamental to ensure the taxonomic rigour of DNA sampling in phylogenetic studies. On a superficial analysis of the first 60 open-access papers on plant phylogenomics retrieved for 2022 on the Google Scholar database (https://scholar.google.com, accessed 10th January 2023), we gathered that only 22% of these studies included a taxonomic expert on the group (i.e., someone who has already published floras or taxonomic revisions in the analysed groups) in the authorship of the paper. Additionally, 78% of these studies do not specify the taxonomic criteria used in the study or mention the taxonomic specialist who would have confirmed the identification of the sampled specimens (see supplementary files). This scenario is the more worrisome since it might be a reflection of the view of biological collections by some plant molecular systematists as immutable DNA archives. This gross misconception of the dynamic nature of plant systematics and the need to constantly revise the determination of the consulted specimens in the light of new evidence can be easily tackled by the association and collaboration with a taxonomic expert in the study group.

While large phylogenomic projects, such as RBG Kew Tree of Life (Baker et al. 2022), have developed thorough quality control pipelines to test the correct phylogenomic placement of the sequenced samples at the family rank (Baker et al. 2022), determinations below the family rank need to be treated with the same care and scrutiny. Furthermore, it is essential to keep good taxonomic practices for collecting the DNA sample itself, as it is not always possible to rely on bioinformatic pipelines to effectively point out the introduced errors in the analyses. Good knowledge of plant taxonomy and morphology is a fundamental basis for minimising mistakes in the early stages of the molecular phylogenetic processes. To make sure that future studies are aware of the dangers of cross-contamination in herbarium specimens or misidentification/contamination in molecular systematic studies, we suggest two sets of golden rules for herbarium sampling:

### Avoiding cross-contamination on herbarium specimens

It is not at all unusual for more than one species to be represented on a herbarium sheet that is labelled as being a single species. Here is how it can happen:

1. *Mixed collections* – In the wild, where species are sampled, two or more species can grow side-by-side or entwined to each other, especially in the case of climbers. If the collector of the specimen is under pressure or is unobservant, specimens that are a mixture can go into the press (commonly known as a “mixed collection” by plant taxonomists).
2. *Look-alike species* – Similar-looking herbaceous species can grow together in close proximity (e.g., species of Gramineae/Poaceae). A collector who needs to fill their press and is not a specialist in the group, and is working under pressure or working under adverse conditions (e.g., poor light or weather), can collect multiple individuals to complete a sheet missing that they are not all the same species, genus, or even from the same family. This is fairly common for certain plant groups, like Monocots, aquatic plants, and small-sized species in general.
3. *Incomplete specimens* – In certain species, inflorescences arise on a plant physically distant from the leaves (e.g., in cauliflorous species, subshrubs, geophytic herbs, aquatic plants, parasitic and mycoheterotrophic species, etc.). A collector can mistakenly associate two species together as a specimen, thinking they belong together. The error may not be detected before the specimen is incorporated into a herbarium and checked by an expert in that group.

### Avoiding misidentification and contamination in molecular plant systematics

In an era of expansion of molecular plant systematic techniques, we here draw attention to the importance of taxonomic skills in guaranteeing the correct sampling for molecular phylogenetic studies and the ability to critically interpret the hypotheses in the light of additional biological evidence before evoking potentially inaccurate results. For future phylogenetic and/or phylogenomic studies sampling herbarium specimens, we suggest simple recommendations to ensure rigorous taxonomic standards to minimise human error and/or taxonomic bias of any kind:

1. *Doubtful identification* – avoid sampling specimens with doubtful identification, i.e., specimens not confidently identified by a taxonomic expert in the plant group (for example, someone who has already published a number of floras or taxonomic revisions). If you are not sure who identified it, consider it doubtful, and keep open the possibility of revisiting the specimen identification, e.g., if the molecular phylogenetic results are to some extent incongruent or inexplicable in the light of the available knowledge about the plant group
2. *Contamination/Misidentification/Mixed specimens* – carefully check the sampled specimens for any mixed collections or contaminations on the mounted sheets, especially inside the capsule storing loose fragments. Loose plant materials are the most prone to contamination and/or misidentification in herbaria, and fragment capsules are always a potential host of mixed leaf material. Make sure the leaves on the capsule are identical to the ones on the mounted specimen, including checking under the stereomicroscope if in doubt.
3. *Nomenclature* – always check on robust online taxonomic databases (e.g., Plant of the World Online, Tropicos.org, etc.) if the name on the identification slip or label is currently accepted by the taxonomic community. This is a fundamental step, very commonly skipped by phylogenetic/phylogenomic studies, that might make it hard for anyone to interpret any taxonomic information in an evolutionary context. In addition to the genus and epithet, also check the authority [e.g., *Ipomoea diversifolia* R.Br. is an accepted name, while *Ipomoea diversifolia* (Schumach. & Thonn.) Didr. is a synonym of *Ipomoea sagittifolia* Burm. f., which is a very different species from *I. diversifolia* R.Br.]. Overlooking this detail, or annotating the species name wrongly, can lead to further complications in the interpretation of a molecular phylogeny, with implications to the systematics of the group in question.
4. *Taxonomist* – the sampler of the phylogenetic/phylogenomic study should, ideally, be someone with good taxonomic experience or willing to receive taxonomic training to prepare them for more easily spotting any incongruencies in the sampled specimens in terms of contamination, material mixture, nomenclature inconsistencies and taxonomic authorship of the determinations on the herbarium samples. Most herbaria rely upon the work of countless past-present-future plant taxonomists to ensure the best taxonomic rigour in curating their collections. Not all plant collections, for several reasons, will have all their specimens always kept up to date according to recent advances in taxonomy and systematics, especially in an age of quick advances in molecular systematics, aggravated by steeping reductions in curatorial and taxonomic staff in most herbaria
5. *Critical thinking* – challenging all preconceptions is vital to the advance of science and must always be the most important part of your study design. Keep an open mind to different explanations as to why results are not as expected, and explore a range of hypotheses. Play detective and retrace your steps, as well as the botanical history of the specimens, to secure that enough evidence, in addition to the molecular data, will support your results.

## Conclusions

Molecular DNA sequences can be very helpful in classifying plant taxa when morphology is conflicting or of a doubtful interpretation, with molecular phylogenetic placement becoming a popular tool to potentially accelerate the discovery of systematic relationships. Nonetheless, it needs to be done with a critical assessment of the obtained results in the context of a range of biological information (i.e., macromorphology, micromorphology, ecology, reproductive biology, phytochemistry, etc.), particularly when the new hypotheses are disruptive to the current classification system. Genetic and genomic techniques are, much like any others, prone to lapses, which further stresses the need for caution in adopting molecular phylogenetic results into a currently accepted classification system. In an era of expansion of molecular plant systematic techniques, we here draw attention to the vital role of morphology and experienced taxonomic skills in guaranteeing an adequate and reliable sampling for molecular phylogenetic studies and the ability to critically interpret the obtained hypotheses in the light of a range of biological evidence, particularly the most easily accessible, morphological characters.

## Acknowledgements

We thank the researchers and curators of all consulted herbaria (Cassio van den Berg, Maria Candida H. Mamede, Maria Lenise Guedes and Viviane Jono) for their assistance; and Climbiê F. Hall, Domingos Cardoso, Fabián Michelangeli, Geovane S. Siqueira, Marco O. O. Pellegrini, Melissa Hutchison, and Olivier Lechenauld for allowing us to use their beautiful photographs. RFA was supported by a postdoctoral fellowship from CNPq (#317720/2021-0) and FAPEG (#202110267000867), Brazil.

